# Changes in attractor dynamics predict altered perceptual decision making with dorsolateral prefrontal tDCS

**DOI:** 10.1101/036905

**Authors:** James J. Bonaiuto, Archy de Berker, Sven Bestmann

## Abstract

The left dorsolateral prefrontal cortex (dlPFC) has been linked to the accumulation and comparison of perceptual evidence for decision making independent of sensory and response modalities. We investigated the possible neural dynamics underlying the role of dlPFC in perceptual decision making, through a combination of noninvasive neurostimulation in humans and computational modeling. First, we used an established and biophysically realistic model of a decision making network that employs competition between neural populations. Simulation of depolarizing noninvasive brain stimulation in this model decreased decision time, while hyperpolarizing stimulation increased it. This behavioral effect was caused by an increase in the rate of neural activity integration via recurrent connections, as well as changes in the susceptibility of the network to noisy background inputs which modulated population firing rate differences prior to the onset of the stimulus. These pre-stimulus differences biased the response to one or the other option, thus speeding or slowing decisions.

We then tested these model predictions in healthy participants performing a perceptual decision making task while receiving transcranial direct current stimulation (tDCS) over the left dlPFC, analogous to our simulated network stimulation. We found a striking match between model predictions and experimental results: depolarizing (inward) currents reduced and hyperpolarizing (outward) currents increased response times, but accuracy remained unaffected. Our results provide interventional evidence for the role of left dlPFC in perceptual decision making, and suggest that this region integrates and compares sensory evidence through competitive interactions between pyramidal cell populations which are selective for each response option. Mechanistically, our model suggests that stimulation of this region changes the rate at which evidence can be accumulated through recurrent activity and its susceptibility to background noise. More generally, our approach demonstrates that a linkage between computational modeling and noninvasive brain stimulation allows mechanistic accounts of brain function to be causally tested.

## Introduction

Perceptual decision making involves comparing sensory information for each option and linking the decision to the appropriate motor responses to indicate the choice. The dorsolateral prefrontal cortex (dlPFC) is thought to be a crucial node in a network of brain regions for categorization and accumulation of continuous perceptual information in decision making [1-11]. In humans, perceptual decision making independent of stimulus and response modality has been attributed to activity in the left dlPFC [12,13]. Interventional evidence with transcranial magnetic stimulation (TMS) of this region shows that stimulation alters the rate at which sensory evidence is accumulated, thus suggesting that this region is indeed necessary for accurate decisions to be made [14]. One way in which dlPFC might accomplish this role is through a competitive process between neural populations which are selective for different choice categories, receive converging input from neurons tuned to values of the continuous perceptual information, and integrate these inputs via recurrent connectivity. The mechanism behind this competition can be described by biophysical attractor models - so called because the network settles into one of several discrete states, or attractors, following input. Such models have been able to explain both choice behavior and the underlying neural firing rate changes in related brain regions [5,7,15,16], as well as population measures of the dlPFC made using functional magnetic resonance imaging (fMRI, [17]) and magnetoencephalography (MEG, [18]). dlPFC dysfunction is common to many neuropsychiatric disorders (e.g. [19-21]); understanding and manipulating dlPFC function is thus of considerable scientific and clinical interest.

One way this has been addressed is through noninvasive brain stimulation (NIBS). This approach seeks to probe or manipulate dlPFC function, but the results have been varied [22-26] and a mechanistic explanation for its effects is currently lacking. Specifically, there is a scarcity of interventional evidence in humans that addresses how alteration of the dynamics in competitive networks within dlPFC may affect perceptual decision making. One notable exception is a recent study using stimulation over dlPFC with transcranial magnetic stimulation (TMS) which produced behavioral effects concordant with a reduction in the rate of sensory evidence accumulation in a drift diffusion model [14]. In this approach, the disruptive pulsed stimulation delivered by TMS leads to the transient disruption of an area (often conceptualized as virtual lesion [27]) which allows for tests of the causal involvement of a region in the specific task of interest. By contrast, we here employed a different form of noninvasive brain stimulation, transcranial direct current stimulation (tDCS), which instead alters neuronal excitability and spontaneous firing rates in brain networks by subtly polarizing membrane potentials through externally applied electrical currents [28-30]. Because tDCS leads to more subtle subthreshold polarization changes rather than synchronized depolarization within an area, we were able to subtly alter the neural dynamics within the targeted brain region noninvasively and *in-vivo*, instead of disrupting ongoing activity [31-33].

One concern for the use of noninvasive brain stimulation is the lack of mechanistic models that can provide predictions about how this intervention might alter neural processes, and hence the resultant behavior. The vast majority of studies employing noninvasive brain stimulation have consulted conceptual models of the effects of stimulation, but these do not consider how neural circuits might produce the observed behavioral change [34]. We have previously advocated simulation, *in silico*, of the neural effects of noninvasive brain stimulation in computational models that can generate behavioral responses, such as the attractor models introduced above, as a way to address this shortcoming [34]. This *computational neurostimulation* approach [15,34] allows for formulating mechanistic explanations for the effects of tDCS on complex and potentially nonlinear neural network dynamics and the resulting changes in behavior. However, there are currently few examples demonstrating how such an approach can be fruitfully applied for the use of NIBS in understanding human cognition [15,35].

To this end, we simulated the acute effects of tDCS [28,30,36] in a biophysical attractor model of dlPFC during a perceptual decision making task. Indeed, recent computational studies suggest that this model might elucidate the mechanistic engine underpinning the effects of tDCS on decision making behavior [15]. The model contains variables and parameters with well-defined anatomical and physiological substrates [7,16,17,37], allowing for explicit simulation and linkage with the known neurophysiological effects of stimulation. This in turn facilitates inspection of the possible mechanistic cause through which stimulation may alter behavior. Our approach thus extends previous work that has utilized more abstract decision variables such as drift diffusion models for identifying through disruption of dlPFC the putative process crucially supported by dlPFC during perceptual decision making [14], and builds upon recent efforts to interrogate the effects of tDCS in neural networks [28,35,36,38]. In the present study, we compared the behavioral predictions generated by the model to data from an analogous experiment in human participants undergoing tDCS targeting left dlPFC, and used the model to explain the neural causes of the behavioral effects of stimulation.

## Results

### Model Architecture

We used an established spiking neural model of decision making implementing an attractor network [15-17]. This model was initially developed to explain the neural dynamics of perceptual decision making [16] and has been used to investigate the behavioral and neural correlates of a wide variety of decision making tasks at various levels of explanation. It is well suited for computational neurostimulation studies because it is complex enough to simulate network dynamics at the neural level, yet is simple enough to generate population-level (neural and hemodynamic) signals, and the resulting behavior, for comparison with human data [17,18]. The model includes model neurons at a level of detail that allows simulation of tDCS by the addition of extra transmembrane currents with parameter values based on previous modeling work [15,36], and current understanding of the mechanism of action of tDCS [28-30,39-41].

The model consists of two populations of pyramidal cells representing the available response options, which are “left” and “right” in this task (Figure 1A). Each population receives task-related inputs signaling the perceived evidence for each response option. The difference between the inputs varies inversely with the difficulty of the task (Figure 1B), and the rate of each input is sampled according to refresh rate of monitor used in our experiment (60Hz, Figure 1C). The pyramidal populations are reciprocally connected and mutually inhibit each other via a common pool of inhibitory interneurons. This pattern of connectivity gives rise to winner-take-all behavior in which the firing rate of one pyramidal population (typically the one receiving the strongest inputs) increases and that of the other is suppressed, indicating the decision. In low coherence trials each input fires at approximately the same rate, while in high coherence trials one input fires at a high rate while the other fires at a very low rate. To simulate the impact of tDCS, we injected an additional trans-membrane current into pyramidal cells and inhibitory interneurons, with the polarity and magnitude based on simulations reproducing tDCS-induced changes in sensory evoked potentials [36] and behavior [15] *in vivo*, and taking into account the cellular effects of tDCS [28-30,39-41].

**Figure 1.**
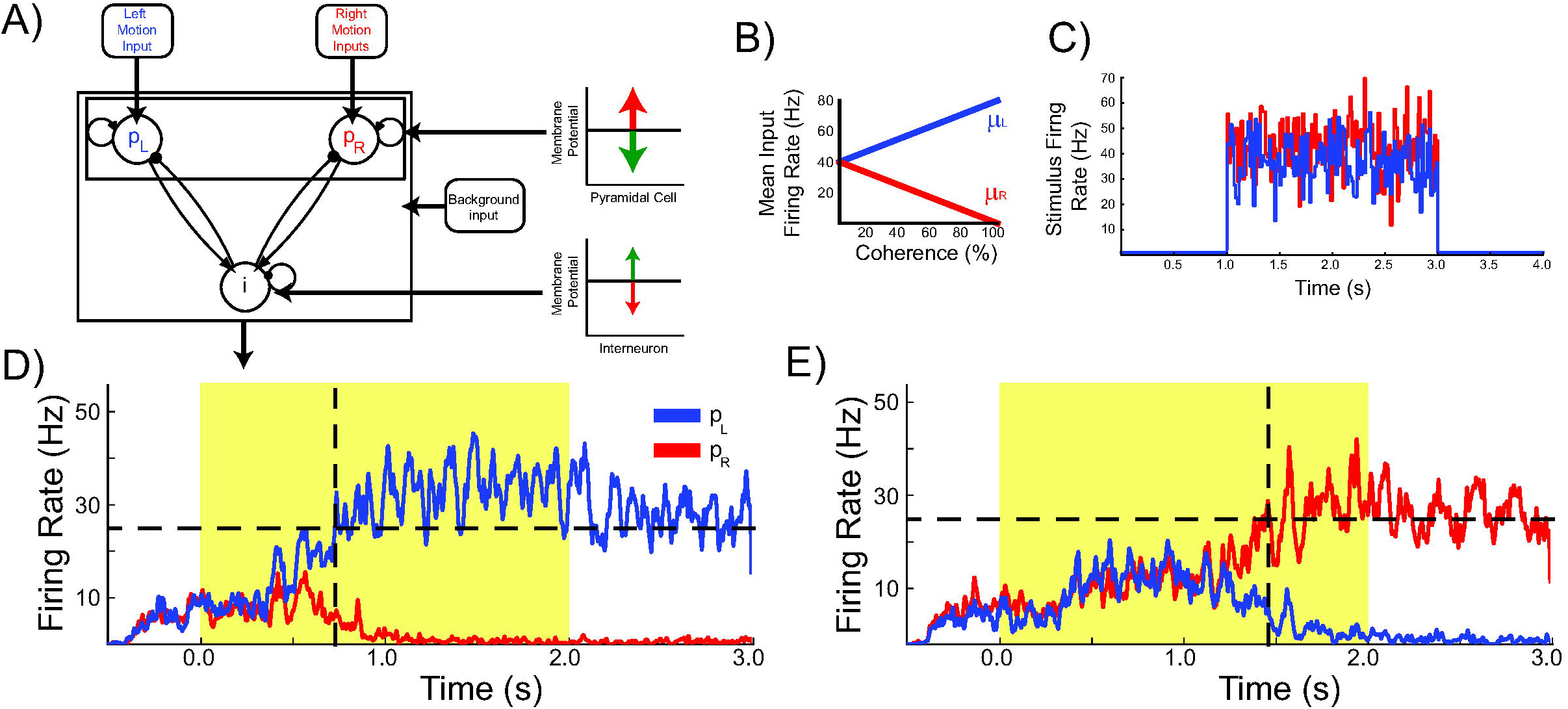
Model architecture, input, and activity. A) The model contains two populations of pyramidal cells which inhibit each other through a common pool of inhibitory interneurons. The pyramidal populations receive task-related inputs signaling the momentary evidence for each response option. tDCS is simulated by modulating the membrane potential of the pyramidal and interneuron populations. B) The mean input firing rate to each pyramidal population, *f.i_L_* and *μ_κ_*, vary as a function of the stimulus coherence. C) The input firing rates are normally distributed and changed according to the stimulus refresh rate of the monitor used in this experiment, to reflect the timing of evidence presentation. D-E) Firing rates of the two pyramidal populations during representative high (D) and low (E) coherence trials. The horizontal dotted lines denote the response threshold (25Hz) and the vertical dotted lines show the decision time - when the first pyramidal population firing rate crosses the response threshold. The decision time is shorter in high coherence trials because the response threshold is crossed earlier.

### Behavioral Results of dlPFC stimulation in silico

We simulated the impact of tDCS on behavior in a perceptual decision making task, by emulating input from a virtual random dot kinetogram (RDK) with varying levels of coherent motion used to scale the magnitude of the task-related inputs (see Methods). The behavior was produced by virtual subjects, which were created by instantiating the model with random fine grained connectivity sampled from distributions defined by its connectivity probabilities. In order to analyze the behavioral output of the network, we consider a response option to be *chosen* when the corresponding pyramidal population exceeds a set *response threshold* of 25Hz. We measured the accuracy of the model’s performance as the percentage of trials in which the chosen option corresponded to the stronger task-related input. For comparison between virtual human participants, we defined the *accuracy threshold* as the coherence level required to attain 80% accuracy. The effects of stimulation on accuracy were determined by comparing the accuracy threshold in each stimulation condition. The time step at which the response threshold is exceeded is taken as the *decision time* for that trial. Because we do not simulate perceptual and motor processes involved in encoding visual stimuli and producing a movement to indicate the decision, this is distinct from the *response time* measured in human participants. The decision time during stimulation conditions were compared to sham no stimulation at each motion coherence level.

As expected, the model generates increasingly accurate responses at higher coherence levels (Figure 2A) because the ‘correct’ pyramidal population is receiving much stronger input than the other, allowing it to more easily win the competition. However, relative to no stimulation, there was no effect of depolarizing or hyperpolarizing stimulation on the model’s accuracy threshold (F(2,38)=1.597, *p*=0.216, *η^2^_ρ_* =0.078; Figure 2A, Table S1). In line with previous work, the model predicts a decrease in decision time with increasing coherence (Wang, 2002; Figure 2D). In terms of model dynamics, when motion coherence is low the sensory evidence for left and right choices is approximately equal, and therefore the inputs that drive both pyramidal populations are more balanced. As a consequence it takes longer for one population to ‘win’ over the other and for the network to reach a stable state (see Figure 1E). For decision time, we observed a main effect of stimulation condition (F(2,38)=136.264, *p*<0.001, *η^2^_ρ_* =0.878) and coherence (F(2.792,53.044)=354.571, *p*<0.001, *η^2^_ρ_* =0.949, ε=0.465) and a significant interaction between stimulation condition and coherence (F(6.176,117.346)=6.702, *p*<0.001, *η^2^_ρ_* =0.261, ε=0.515, Table S2). The model thus predicts increased accuracy and faster responses with increasing motion coherence. Moreover, it predicts that network stimulation will affect response time, but leave accuracy largely unaffected. It is predicted that depolarizing and hyperpolarizing stimulation will lead to faster and slower responses, respectively.

**Figure 2.**
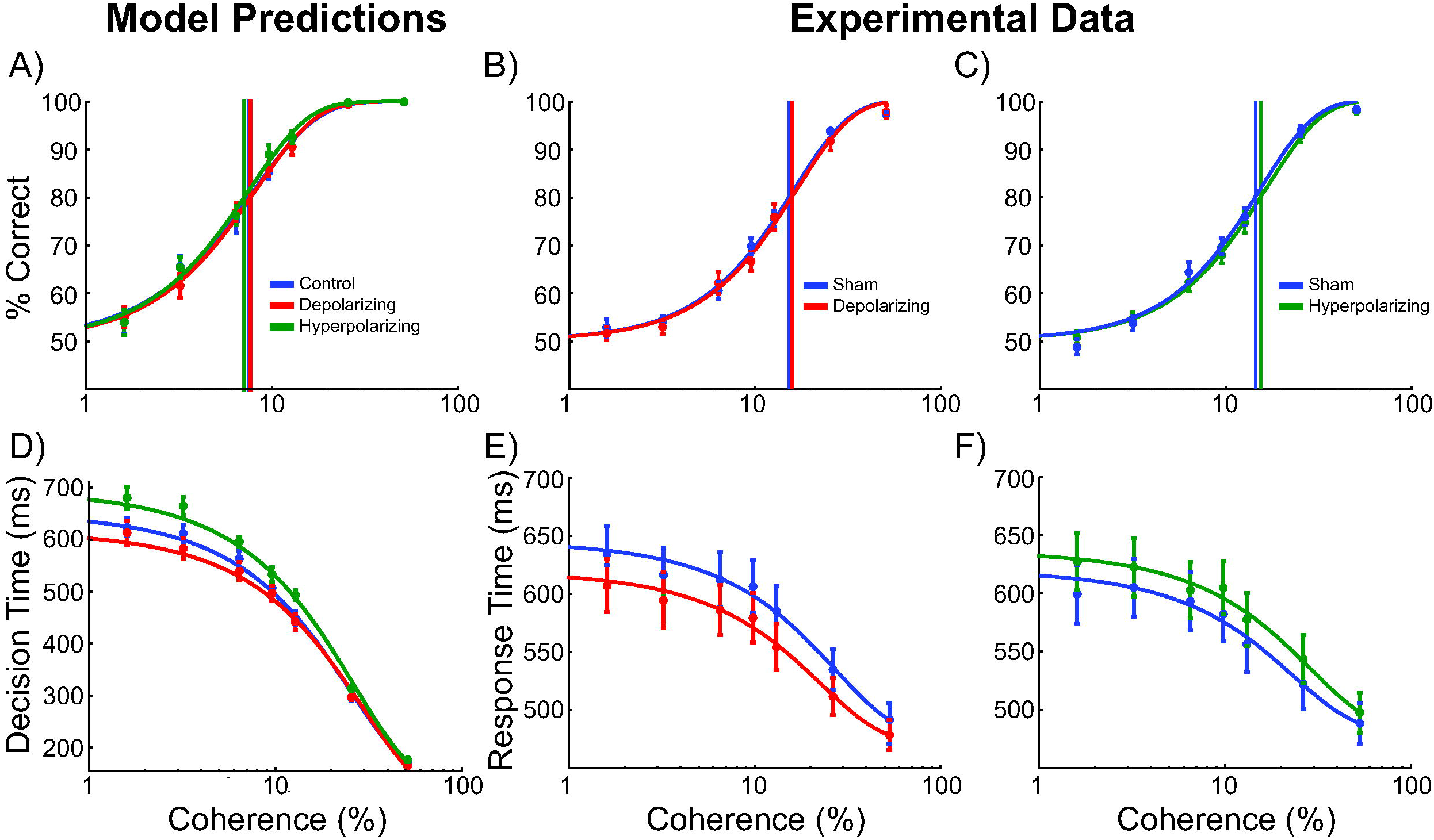
Behavioral effects of stimulation. A) The model predicted no change in the decision threshold - the coherence required to reach 80% accuracy with stimulation. B-C) Neither depolarizing nor hyperpolarizing stimulation altered the decision thresholds of human participants. D) The model predicted a decrease in decision time with depolarizing stimulation and an increase with hyperpolarizing stimulation. E-F) In human participants, depolarizing stimulation decreased and hyperpolarizing stimulation increased decision time relative to sham stimulation.

As discussed above, the model predicts a reduction of decision time during depolarizing stimulation and an increase during hyperpolarizing stimulation. In order to investigate the interaction between stimulation and task difficulty on model decision time, we analyzed the difference in decision time between sham and stimulation at each motion coherence level. In both stimulation conditions, this difference gradually disappears with increasing coherence as indicated by the significant interaction between stimulation condition and coherence (F(2.607,49.526)=49.241, *p*<0.001, *η^2^_ρ_* =0.722, ε=0.434; Figure 3A, Table S3). This is because during trials with high coherence, the network is already near the lower bound of the decision time, which is determined by neural time constants and population dynamics. The external perturbations induced by stimulation do not overcome the stable attractor state that the network quickly reaches under such circumstances. These simulations therefore predict that response time should be unaffected on ‘no brainer’ trials in which strong inputs provide unequivocal evidence for one response over the other. This echoes findings that tDCS may interact with task difficulty and/or individual differences in performance [42,43].

**Figure 3.**
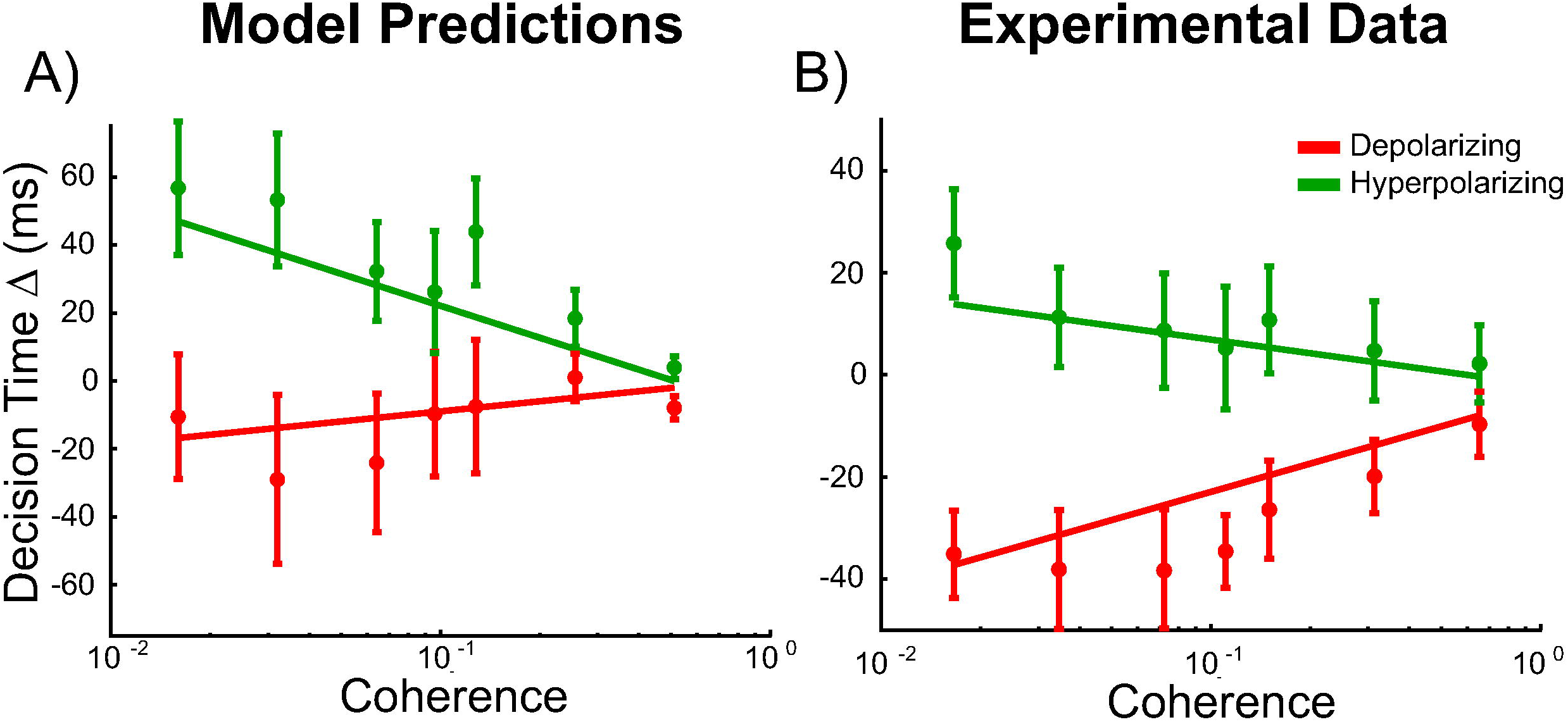
Effects of stimulation on decision time. A) The model predicts that while depolarizing stimulation decreases and hyperpolarizing stimulation increases decision time, the effect is reduced with increasing coherence. B) In human participants depolarizing and hyperpolarizing stimulation significantly decreased and increased the decision time, respectively, matching the model predictions shown in A.

We next asked whether the predictions from our simulated stimulation were borne out in the behavior of real subjects undergoing tDCS.

### Behavioral Results of dlPFC stimulation in vivo

24 human participants performed the same perceptual decision making task simulated in the model, in which they viewed a RDK and were required to indicate the direction of coherent motion (Figure 4A). The accuracy of the human participant performance was measured as the percentage of trials in which participants correctly indicated the direction of coherent motion, and each participant’s accuracy threshold was defined as the motion coherence required to reach 80% accuracy. Four of these participants were excluded from the analysis due to poor performance on the task (see Methods). Transcranial Direct Current Stimulation (tDCS) was applied over the left dorsolateral prefrontal cortex (Figure 4B) in order to induce depolarizing, hyperpolarizing, or sham stimulation. Stimulation blocks were conducted in separate sessions, to avoid carry-over effects of repeated stimulation blocks, and compared with the directly preceding sham block in the same session (Figure 4C). We therefore have separate baselines for each stimulation condition, and consequently separate plots for accuracy and response time in Figure 2.

**Figure 4.**
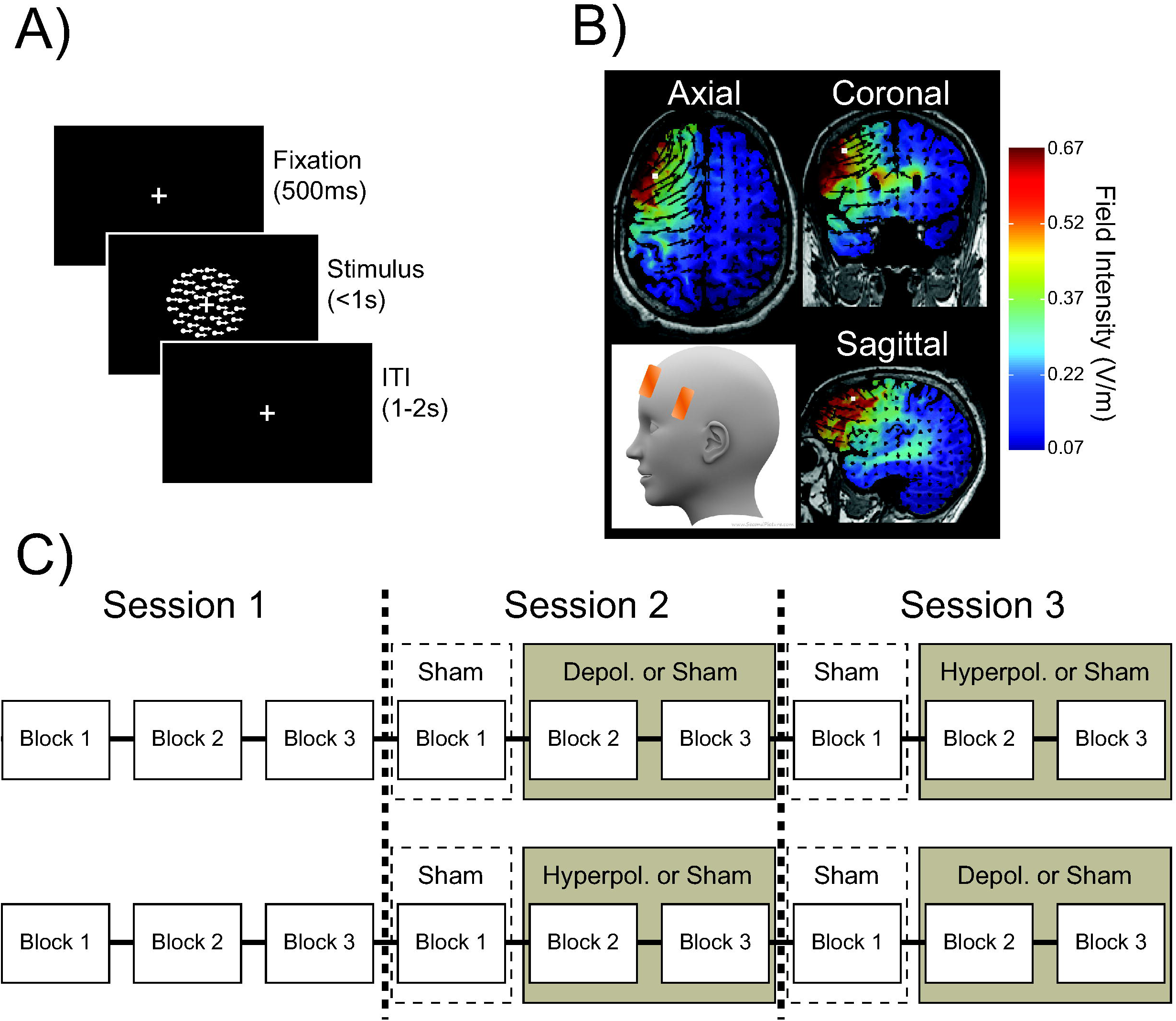
Behavioral task, electrode position and current stimulation, and session order. A) Behavioral task B) Electrode positions and estimated current distribution during left dorsolateral prefrontal stimulation. C) Experimental protocol.

As predicted by the model and in line with previous work [44], the accuracy of human participants increased (Figure 2B-C) and response times decreased (Figure 2E-F) with increasing motion coherence. In striking accordance with the predictions of our biophysical model, neither depolarizing nor hyperpolarizing stimulation had an effect on the accuracy threshold, but tDCS over left dlPFC altered response time.

Compared to sham stimulation, the accuracy threshold of human participants was not changed by depolarizing (t(20)=0.523, *p*=0.607, d=0.131) or hyperpolarizing (t(20)=0.588, *p*=0.563, d=0.132) stimulation (Figure 2B). However, as predicted, there were main effects of stimulation condition (F(3,60)=4.081, *p*=0.011, *η^2^_ρ_* =0.169) and coherence (F(6,120)=78.310, p<0.001, *η^2^_ρ_* =0.797) on decision time, as well as an interaction between stimulation condition and coherence (F(18,360)=1.686, *p*=0.04, *η^2^_ρ_* =0.078) on the response time of human participants (Figure 2E-F).

To test the model prediction concerning the interaction between stimulation and task difficulty on decision time, we analyzed the response time of human participants in each stimulation block, compared to the preceding sham block at each motion coherence level after smoothed them using a moving average with a window size of 3. Indeed, and exactly as predicted by the model, human participants showed the same pattern of effects on relative response time for depolarizing and hyperpolarizing stimulation compared to sham stimulation (condition x coherence interaction: F(2.545,50.891)=8.18, *p*<0.001, *η^2^_ρ_* =0.29, ε=0.424, Figure 3B).

### Neural Effects of Simulated tDCS

We found that in both our model and in human participants, polarization of the dlPFC led to changes in reaction times in a perceptual decision making task, in line with previous work assigning an important role to dlPFC in categorization and accumulation of perceptual evidence [2,8,14]. One advantage of combining experimental human studies with computational models is that it allows for interrogation of the neural dynamics of the model under different experimental manipulations [45,46]. We therefore asked how the injection of current into our model altered its dynamics, producing the changes in model behavior that we observed. This ability to ‘get under the hood’ of biophysical models to illustrate the processes underlying stimulation effects is a unique property of computational neurostimulation approaches, and in the absence of invasive recordings can help to generate mechanistically grounded hypotheses on how stimulation of a neural network produces behavioral change.

As can be seen in Figure 5A, each pyramidal population fires at approximately 5Hz prior to the onset of the task-related inputs. These inputs initially increase the firing rate in both pyramidal populations, which drive increased activity in the interneuron population. We sorted the population firing rates of each trial based on which pyramidal population was eventually chosen. We found that the mean firing rate of the chosen population was slightly higher than that of the unchosen population prior to onset of task-related input. Because neurons from each population receive background (non-task-related) input at the same rate, this effect can be attributed to random fluctuations in the total amount of noisy background input. We refer to this difference in firing rate prior to the onset of the task-related input as *pre-stimulus bias*. Because stimulation was applied throughout the entire simulation and not just for the duration of the task-related stimuli, this matched the actual application of tDCS in the behavioral experiment.

**Figure 5.**
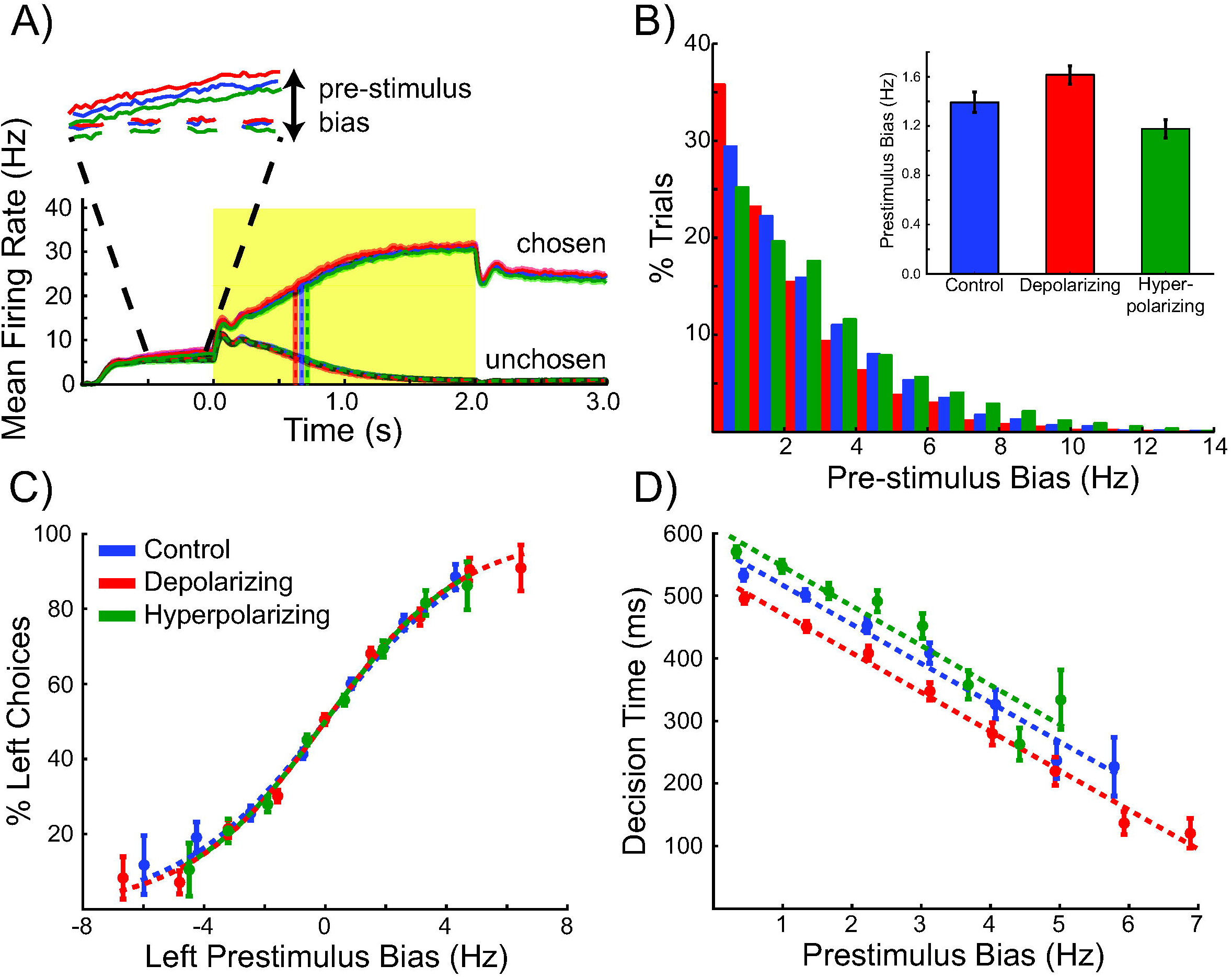
Pre-stimulus bias accounts for the effect of stimulation upon reaction time. A) When sorted according to chosen and unchosen categories, stimulation increased the difference between the firing rates of the chosen (solid lines) and unchosen (dashed lines) pyramidal populations prior to the onset of the task-related stimulus. B) Depolarizing stimulation increases pre-stimulus bias while hyperpolarizing stimulation decreases it (mean and standard error shown in the inset). C) Stimulation slightly changes the range of pre-stimulus bias values, but the effect on the selection remains constant across conditions. D) Decision time and pre-stimulus bias are negatively correlated, showing that being biased to one option reduces the time the biased pyramidal population takes to reach the threshold firing rate. Stimulation reduces (depolarizing) or increases (hyperpolarizing) decision time, but the slope of the relationship between the pre-stimulus bias and decision time is the same.

Depolarizing network stimulation effectively amplifies pre-stimulus bias, whereas hyperpolarizing stimulation suppresses it (F(2,38)=77.174,*p*<0.001, *η^2^_ρ_* =0.802, Figure 5B, Table S4). All pyramidal neurons received the same level of stimulation, thus stochastic fluctuations in the differences in background input between the two populations were amplified or suppressed.

Having demonstrated that depolarizing and hyperpolarizing stimulation differentially affect the pre-stimulus bias, we next asked how pre-stimulus bias affected accuracy in our model. Large pre-stimulus biases influence network dynamics by endowing the pyramidal population with the higher initial firing rate with a head-start. A large enough pre-stimulus bias would make it much more likely to win the competition, irrespective of whether it subsequently receives the stronger task-related input [15]. The relationship between the pre-stimulus bias and the percent of trials in which the biased population wins is well described by a sigmoid function (Figure 5C), but the steepness parameter is not changed by stimulation (F(2,38)=1.129, *p*=0.334, *η^2^_ρ_* =0.056, Table S5). Therefore while stimulation affects the *mean* pre-stimulus bias (Figure 5B), the effect of the pre-stimulus bias itself on the choices made by the model is not changed. In order to significantly affect choice accuracy, stimulation must change the mean pre-stimulus bias by a large enough amount to shift the dynamics closer to the extreme ends of the sigmoid shown in Figure 5B. Under the stimulation intensities tested here, stimulation did not change the range of the pre-stimulus bias enough to significantly alter choice behavior (and thereby accuracy, see Discussion). However, we note that stimulation at higher intensities can potentially affect accuracy, although such intensities are currently not commonly applied in human tDCS studies [15].

Although the model predicts no effect of stimulation on accuracy, decision time was decreased and increased by depolarizing and hyperpolarizing stimulation, respectively. We therefore asked how pre-stimulus bias affects model decision time. Large pre-stimulus biases lift one pyramidal population slightly closer to the response threshold, reducing the time needed to reach that threshold. This would reduce decision time in trials in which that population eventually wins, but increase decision time in trials in which the other population receives task-related inputs strong enough to overcome the bias. However, there was a negative linear relationship between the pre-stimulus bias and decision time for all stimulation conditions (Figure 5D). The offset of this relationship is the mean decision time when the pre-stimulus bias is zero. The slope describes the change in decision time as the head start given to one of the pyramidal populations by the pre-stimulus bias increases. The slope of this function is not affected by stimulation condition (F(1.·497,28.·442)= 1.05, *p*=0.344, *η^2^_ρ_* =0.052, ε=0.748, Table S6), showing that similar to accuracy, the stimulation does not change the head start effect of pre-stimulus bias on decision time. However, the difference in the offset induced by stimulation (F(2,38)=9.012, *p*=0.001, *η^2^_ρ_* =0.322, Table S6) demonstrates that the effects of stimulation on decision time, unlike accuracy, extend beyond the pre-stimulus period (stimulation is applied throughout the duration of the simulation).

This observation can be explained by the variation in the decision time with the stimulation condition for the same level of pre-stimulus bias. Therefore, the differences in decision time are not due to the effects of stimulation in the pre-stimulus period. In other words, even with equivalent levels of pre-stimulus bias, the mean decision time is shorter for depolarizing stimulation and longer for hyperpolarizing stimulation: even after the onset of the task-related input, the stimulation is amplifying (depolarizing) or dampening (hyperpolarizing) the effects of the bias. However, the pre-stimulus bias can be either congruent or incongruent with the subsequent task-related inputs. In each case, the resultant decision can be a correct or incorrect. When pre-stimulus bias was congruent with the input, correct responses (consistent with the bias) were made over a wide range of bias values, while incorrect responses were only made when the pre-stimulus bias was less than 3Hz (Figure 6A). This means that ongoing noisy background inputs only reversed the (correct) pre-stimulus bias and induce an incorrect response when the bias was very small. When the pre-stimulus bias was incongruent with the task-related inputs, incorrect responses (consistent with the bias) were made over a wider range of prestimulus bias values than correct responses, indicating that a larger bias is required to overcome the task-related inputs (Figure 6D).

**Figure 6.**
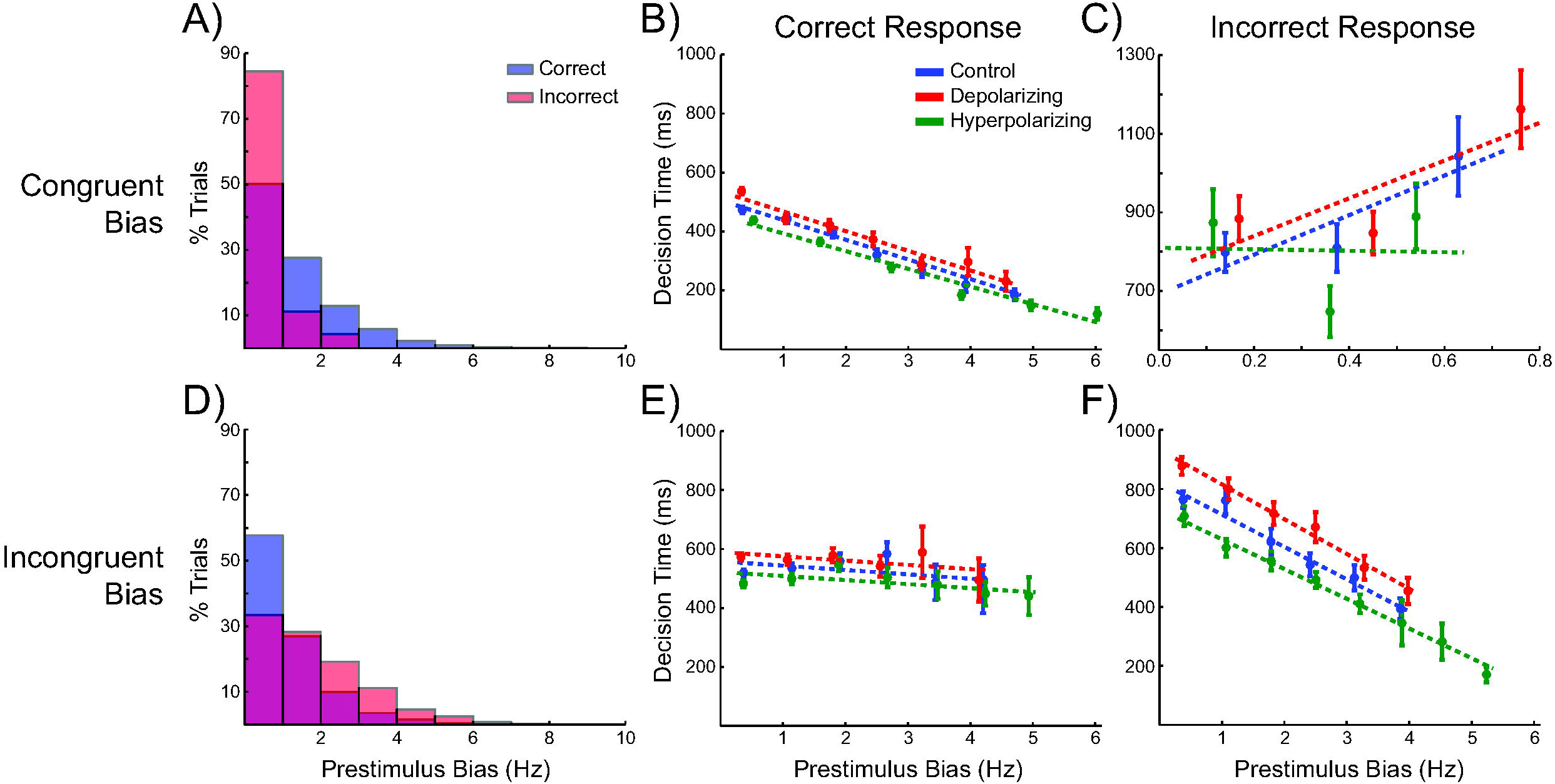
The congruence between the pre-stimulus bias and task-related inputs explains the effect of pre-stimulus bias upon reaction time. A) When the pre-stimulus bias is congruent with the subsequent task-related inputs, correct responses are made over a wide range of pre-stimulus bias values, while incorrect responses are only made when the bias is low. B) When a correct response is made during congruent trials, the decision time decreases with increasing bias. C) In the few congruent trials in which an incorrect response is made, the decision time was much longer and there was no consistent effect of stimulation. D) When the pre-stimulus bias is incongruent with the subsequent task-related inputs, incorrect responses are made over a wider range of pre-stimulus bias values than correct responses. E) When a correct response is made in spite of an incongruent pre-stimulus bias, the task-related inputs quickly overcome the bias, resulting in no effect of pre-stimulus bias on decision time. F) The largest effect of the pre-stimulus bias on decision time occurs during incongruent trials in which an incorrect response is made.

How do these differences explain the effects of stimulation on decision time? We found that the largest effect of pre-stimulus bias on decision time and the largest variation between conditions occurred when the bias was incongruent with the task-related inputs and was large enough to overcome them and induce an incorrect response (Figure 6F). In trials in which the bias was congruent with the task-related inputs and a correct response was made, there was a small effect of bias value on decision time (Figure 6B). In the very few trials where the bias was congruent with the task-related inputs but an incorrect response was made (mostly very difficult low coherence trials), it took a long time to overcome the bias because of the small difference in task-related inputs and there was no consistent relationship between the pre-stimulus bias level and decision time (Figure 6C, note the difference in scale). Finally, when the bias was incongruent with the inputs but was overcome in order generate a correct response, the prestimulus bias provided a head start to the wrong population, but this was quickly overcome and resulted in no effect of pre-stimulus bias on decision time (Figure 6E). While the number of trials where the bias is congruent or incongruent with the inputs is roughly similar, there are more correct responses when the pre-stimulus bias is large and incongruent with the task-related inputs (Figure 6D, blue bars) than incorrect responses when it is congruent (Figure 6A, red bars). This is because at this level of stimulation intensity, pre-stimulus bias values tend to be at the lower end of the range in task-related input differences, and the bias can therefore more easily be overcome when it is incongruent with the inputs. This explains why, when all of the trials are averaged together, decision time decreases with increasing pre-stimulus bias (Figure 5D).

## Discussion

The dlPFC is thought to be crucial for perceptual decision making, by integrating and comparing categorized sensory evidence regardless of stimulus and response modality [4,8,12,13]. Mechanistically, it has been proposed that this integration occurs through competitive dynamics in neural circuits containing populations of pyramidal cells tuned to each response option [5,7,16]. The fact that TMS applied over the left dlPFC impairs perceptual decision making suggests that dlPFC indeed plays a causal role for perceptual decision making [14]. However, the putative neural mechanisms and dynamics underlying this role have not been investigated with a similar causal approach.

We here used tDCS, a noninvasive neurostimulation technique, over left dlPFC to provide subtle perturbations of neural network dynamics while participants performed a perceptual decision making task. We leveraged recent developments in computational modeling approaches that bridge between the physiological and behavioral consequences of tDCS [45,47-49] to generate behavioral predictions about how the gentle perturbation of network dynamics might impact behavior. This is also an important step towards understanding the impact of brain stimulation: given the non-trivial and non-linear dynamics that any population-based interference elicits, model-free interpretations of its behavioral consequences likely fail to disclose the likely mechanisms underpinning stimulation effects [34]. To this end, we simulated the impact of tDCS in an established biophysical attractor model of dlPFC function. We show that stimulation not only alters the rate of increase in the firing rates of recurrently connected pyramidal populations, but also modulates firing rate differences prior to the onset of task-related stimuli. This in turn biases choice behavior to one option or the other in the same way we observed in healthy participants undergoing analogous stimulation over dlPFC. In doing so, we validate the biophysical model’s ability to predict the behavioral effects of injected currents, while simultaneously providing a mechanistic account of dlPFC stimulation’s effects upon human behavior during perceptual decisions.

### Behavioral effects of stimulation in silico are matched by analogous stimulation over left dlPFC in healthy participants

In our simulations, stimulation had no discernible effect on choice accuracy, but depolarizing stimulation decreased decision time, while hyperpolarizing stimulation increased it. Moreover, the effect of stimulation on decision time increased with increasing task difficulty. These effects were largely explained by the amplification or suppression of the influence of stochastic background activity on pyramidal cell firing before the onset of the stimulus, owing to the polarizing currents that were applied throughout the task, thereby influencing the prestimulus activity.

We then tested the behavioral predictions made by the model in a group of human participants performing the same task, and found a striking match between their behavioral performance and the model predictions, both in terms of the differential effects on response times and accuracy, but also the polarity of stimulation (depolarizing/hyperpolarizing). These results provide interventional evidence for the role of dlPFC in perceptual decision making [1,2,10-14,50,51], and suggest that this region acts as an attractor network in which perceptual evidence is accumulated and a decision made.

We found that both stimulation conditions significantly affected the model’s decision time, but not choice accuracy. The primary mechanism behind the effect of stimulation on decision time was a modulation of the difference in pyramidal population firing rates prior to the onset of the task-related stimulus, which we refer to as pre-stimulus bias. Even without simulated tDCS, pre-stimulus bias can occur because of random variations in the background (non-task-related) inputs, which all neural populations in the model receive. In our simulations, prestimulus bias was enhanced by depolarizing and suppressed by hyperpolarizing stimulation, therefore effectively increasing (depolarizing) or decreasing (hyperpolarizing) the influence of the non-task-related inputs on the network dynamics. For example, when the pre-stimulus bias is high, choice variability is driven almost entirely by stochastic background inputs, rather than by variability in the task-related inputs, which dominate dynamics when pre-stimulus is low.

### Computational neurostimulation offers a window into the mechanistic engine driving stimulation-induced behavioral change

Pre-stimulus bias influenced decision time by endowing the ‘biased’ pyramidal population with a head-start, by increasing its average firing rate closer to the response threshold. Depolarizing stimulation decreased, and hyperpolarizing stimulation increased decision time, and both had a greater effect on difficult trials with low motion coherence. In addition to the impact of pre-stimulus bias, the effects of stimulation extended throughout the trial - in all conditions, the average decision time linearly decreased with increasing pre-stimulus bias. The slope of this relationship did not vary with stimulation type, meaning that in each condition, the effect of a given amount of pre-stimulus bias compared to no bias was the same. However, the offset was different between conditions, revealing that the decision time varied between conditions even on trials with very small levels of pre-stimulus bias. This indicates that the effects of stimulation on decision time were not solely due to pre-stimulus effects, and that Stimulation increased (depolarizing) or decreased (hyperpolarizing) the speed of the increase in firing rate of the winning pyramidal population.

A surprising prediction of our model that was confirmed with human participants is that stimulation affects decision time but not choice accuracy. In previous studies with a similar model we found that decision making accuracy is affected at higher intensity levels of stimulation [15]. While stimulation can amplify or suppress pre-stimulus bias, which affects choice behavior, the stimulation intensity used in our simulations were not high enough to modulate pre-stimulus bias by an amount sufficient to completely override differences in task-related inputs (left/right motion coherence). In other words, while the network polarization afforded by tDCS modulates network dynamics and influences the speed at which a decision bound is reached, it does not prevent task-related inputs from driving the network, at least not within the plausible intensity levels we used here. This is in contrast with previous work showing that repetitive TMS over left dlPFC reduces accuracy and increases response time [14]. However, whereas TMS elicits instantaneous synchronized activity within the area of stimulation, tDCS subtly alters neural dynamics through de- or hyper-polarizing currents without directly eliciting spikes [40,52]. Consequently, rather than resetting or bluntly disrupting activity within dlPFC, our tDCS intervention altered the neural dynamics without preventing the network from operating. Such a conclusion is compatible with our simulation results, and animal work on the impact of tDCS [15,36,53,54]. The subtle alteration in dynamics rather than disruption may account for the interaction between stimulation condition and coherence level in our study, while previous work using rTMS to the left dlPFC was found to alter performance similarly across task difficulty levels [14]. Another important difference between the two approaches is that TMS applies very short and precisely timed impulses at specific stages within a trial, whereas tDCS here was applied throughout the experiment. This may also account for the observation that stimulation not only altered decision time consistent with a change in the rate of evidence accumulation [14], but also caused pre-stimulus bias effects which would correspond to starting point differences in a drift diffusion model.

### Study Limitations

Here, we simulated the effects of both depolarizing and hyperpolarizing dlPFC stimulation. In these simulations, currents affected both pyramidal cells as well as interneurons. This modeling choice was based on previous work showing that simulated tDCS must affect both pyramidal cells and interneurons in order to explain changes in sensory evoked potentials observed *in vitro* [36]. Some accounts of the neurophysiological effects of tDCS suggest that pyramidal neurons are predominantly affected [40,55], but in additional simulations in which stimulation was only applied to the pyramidal cells the results were qualitatively similar (Table S1-S6).

The level of stimulation intensity we used is based on current modeling estimates of the mean field strength in dlPFC and *in vitro* measurements of pyramidal cell and interneuron polarization as a function of field strength [30]. We used individual structural MRIs to optimize electrode placement for each participant. Electrode positions relative to the MNI template were determined using current modeling to maximize current flow through the superior frontal sulcus portion of the left dlPFC. Each participant’s MRI was aligned to the template and the inverse of this transformation was used to derive optimal electrode positions to generate current flow through the same sulcus. We are therefore confident that our electrode montage accurately targeted left dlPFC and that our simulated tDCS intensity matched the polarization of human neurons using 2mA tDCS. However, in our simulations neurons were not spatially localized and polarization was applied uniformly to all neurons within a population. Future computational neurostimulation studies should investigate the effects of heterogeneous polarization due to variable patterns of current flow through brain tissue.

### Computational Neurostimulation

tDCS is widely used in basic and translational studies for reversible and controlled modulation of neural circuit activity in the human brain. However, there is a distinct lack of mechanistic models that not only explain how stimulation affects neural network dynamics, but also how these changes alter behavior. There are several conceptual models of the effects of noninvasive brain stimulation, but the explanations that they offer typically make leaps across several levels of brain organization and don’t consider how neural circuits generate behavior [34]. Efforts have been made in this direction [28,36,45,48,49,56,57], but there have been very few computational models that offer an explanation for the behavioral effects of tDCS in terms of neural circuit dynamics [15,35]. We here present the first biophysically informed modeling study of tDCS effects during perceptual decision making, and provide detailed hypotheses about the changes in neural circuits that translate to observed behavioral changes during stimulation.

The major advantage of computational neurostimulation is that it uses biophysically informed models which allow for concrete mapping of model dynamics to neurophysiological processes. In addition to making simulation of the effects of tDCS relatively straightforward, this allows model activity to be directly interpreted in terms of neural dynamics and make predictions that easily translate to neurophysiology experiments. Such an approach extends previous approaches for investigating the causal relevance for a brain region in a cognitive process such as perceptual decision making [14], by testing predictions made by mechanistic biophysical models of that region in an interventional way, rather than fitting the output of a behavioral model to human responses and explaining the effects of stimulation in terms of changes in abstract parameter values.

Another beneficial feature of computational neurostimulation is that it furnishes very specific hypotheses for future experiments. For example, our model explains the effects of tDCS on response time through a pre-stimulus bias which modulates differences in pyramidal population firing rates before the onset of the stimulus. This suggests the design of a trial-wise triggered tDCS experiment in which stimulation is only applied before sensory evidence is provided. Our model predicts that response time will be affected proportionally by the prestimulus bias. Sliding-scale conceptual models of the effects of tDCS suggest that it dials up (anodal) or down (cathodal) the functionality in a region, but in a dynamic system it may be difficult to predict how behaviour should be affected by different stimulation protocols without the consultation of a model. In the present study, our model predicts that the pre-stimulus bias and rate of integration will be amplified and suppressed by de- and hyper-polarizing stimulation, respectively.

## Conclusion

We have shown that a biophysical attractor model generates perceptual decision making behavior accurately matching that of human participants. Additionally, we used computational neurostimulation of this model to predict the effect of tDCS over left dlPFC, which we then confirmed experimentally. Previous work showing that changes to parameters in diffusion models can explain differences in human perceptual decision making after transcranial magnetic stimulation of left dlPFC [14]. Our results extend these findings by address the neural circuitry behind perceptual decision making processes, allowing us to offer mechanistic explanations for the effects of stimulation on neural dynamics in the left dlPFC. We provide interventional evidence that the left dlPFC integrates and compares perceptual information through competitive interactions between neural populations which are selective for each response category.

## Methods

### Biophysical Attractor Model

The model contains two populations of pyramidal cells representing the “left” and “right” choice options. These populations of pyramidal neurons *(p_L_* and *p_R_*, 800 neurons each) receive task-related inputs, form excitatory reciprocal connections with neurons in the same population, and mutually inhibit each other via a common pool of 400 reciprocally connected inhibitory interneurons. The neurons in the two pyramidal populations project to excitatory synapses (AMPA and NMDA) on target cells and the interneurons project to inhibitory synapses on their targets (GABA_A_).

All neural populations receive stochastic background input from a common pool of Poisson spike generators, causing each neuron to spontaneously fire at a low rate. We used different background input firing rates sampled from a range previously used to simulate human participants in a similar decision making task [15]. The pyramidal populations additionally receive task-related inputs signaling the perceived evidence for each response options using the same scheme as Wang [16]. The mean rates of the task-related inputs, *μ_L_* and *μ_R_*, vary linearly with the coherence level of the simulated RDK (Figure 1B). Importantly, the sum of the mean task-related input rates always equals 80Hz, meaning that decision making behavior has to emerge from network dynamics and input structure and cannot be attributed to differences in the overall level of task-related input stimulation. The firing rate of each task-related input at each time point was normally distributed around the mean (σ=4Hz) and changed according to refresh rate of monitor used in our experiment (Figure 1C). In additional simulations we show that the behavior of the model is qualitatively robust to changes in the total task-related input firing rates and refresh rate (Table S1-S6).

For analysis, we compute mean population firing rates by convolving the instantaneous population firing rate with a Gaussian filter 5ms wide at the tails. The winner-take-all dynamic of the network causes the firing rates of the pyramidal populations to magnify differences in the inputs. This is due to the reciprocal connectivity and structure of the network, which endows it with bistable attractor states, resulting in competitive dynamics [58,59]. As the firing rate of one population increases, it increasingly inhibits the other population via the common pool of inhibitory interneurons. This further increases the activity of the winning population as the inhibitory activity caused by the other population decreases. These competitive dynamics result in one pyramidal population (typically the one receiving the strongest input) firing at a relatively high rate, while the firing rate of the other population decreases to approximately 0Hz (Figure 1D-E).

### Synapse and Neuron Model

Here we provide a detailed description of the architecture of the biophysical attractor model we used. We modeled synapses as exponential (AMPA, GABA_A_) conductances, or biexponential conductances (NMDA). Synaptic conductances are governed by the following equation:

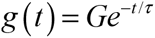

where *G* is the maximal conductance (or weight) of that specific synapse type (AMPA, GABA_a_, or NMDA), and *τ* is the decay time constant for that synapse type. Similarly, bi-exponential synaptic conductances are determined by:

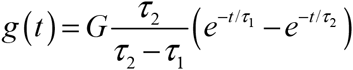

where *τ_2_* and *τ_2_* are decay and rise time constants. Synaptic currents are computed from the product of these conductances and the difference between the membrane potential and the synaptic current reversal potential, *E*:

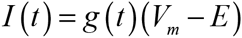

where *V_m_* is the membrane voltage. NMDA synapses have an additional voltage dependence, which is captured by:

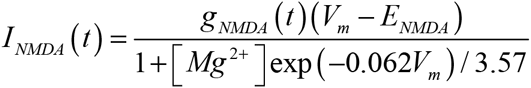

where [Mg^2+^] is the extracellular magnesium concentration.

The total synaptic current (summing AMPA, NMDA, and GABA_A_ currents) is input into the exponential leaky integrate-and-fire (LIF) neural model [60]:

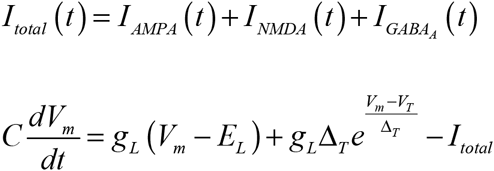

where *C* is the membrane capacitance, *g_L_* is the leak conductance, *E_L_* is the resting potential, *Δ_T_* is the slope factor (which determines the sharpness of the voltage threshold), and *V_T_* is the threshold voltage. After spike generation, the membrane potential is reset to *V_R_* and the neuron cannot generate another spike until the refractory period, *τ_R_*, has passed. Intra- and interpopulation connections are initialized probabilistically with axonal conductance delays of 0.5ms. Parameter values are based on experimental data from the literature where possible [61-65] and set empirically otherwise (Table 1).

All connection probabilities are determined empirically so that the network generates winner-take-all dynamics [15,17]. Recurrent pyramidal population connectivity probability (the probability that any pyramidal cell projected to an AMPA or NMDA synapse on other cells in the same population) was 0.08, and recurrent inhibitory interneuron population connections used GABA_A_ synapses and had a connectivity probability of 0.1. Projections from the pyramidal populations connected to AMPA or NMDA synapses on the inhibitory interneurons with probability 0.1, connections from the inhibitory interneuron population to each pyramidal population used GABA_A_ synapses with a connectivity probability of 0.2. Thus, the pattern of connectivity between populations was fixed, but the fine-scale connectivity between individual neurons was probabilistically determined by the connectivity parameters.

**Table 1.**
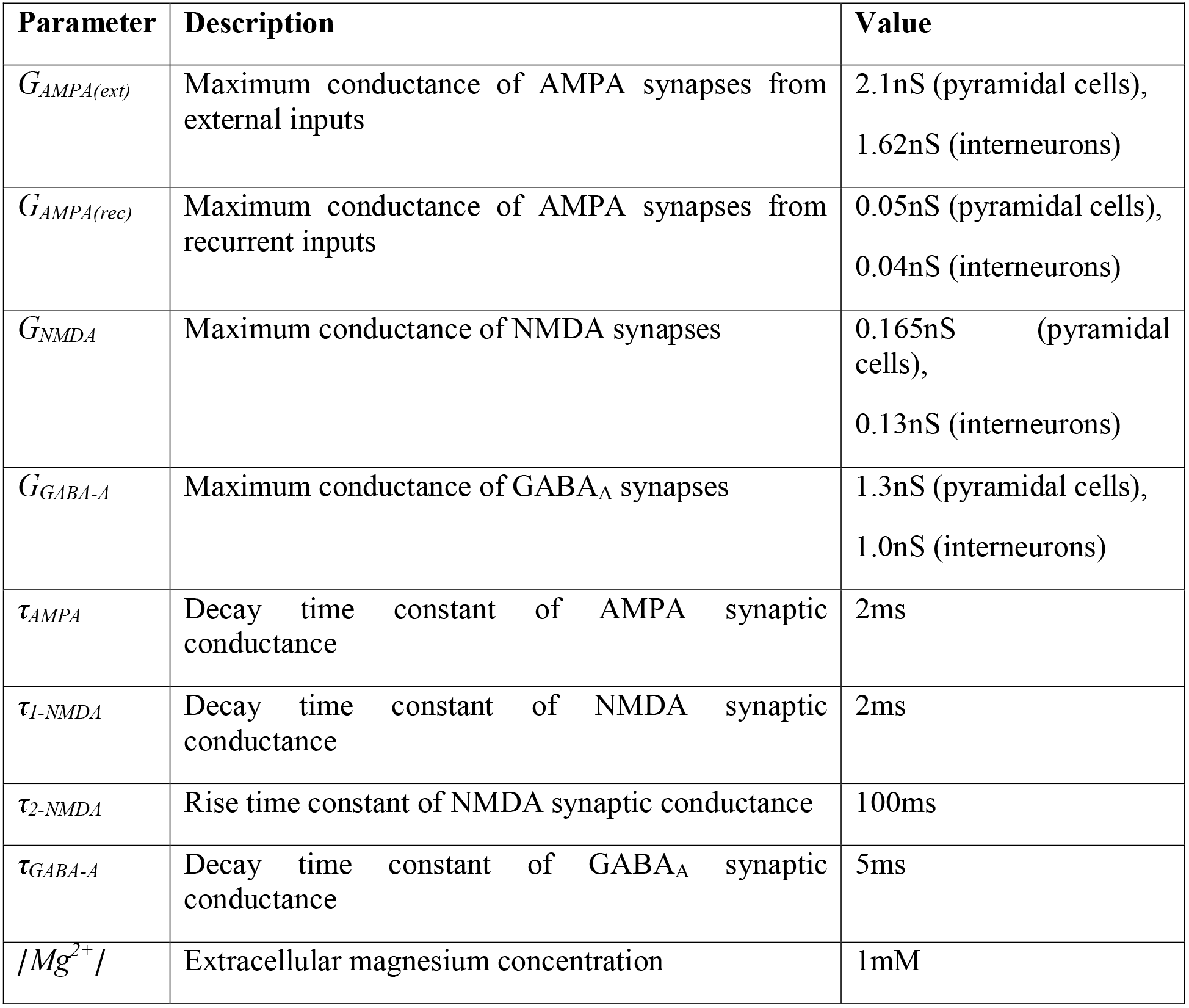
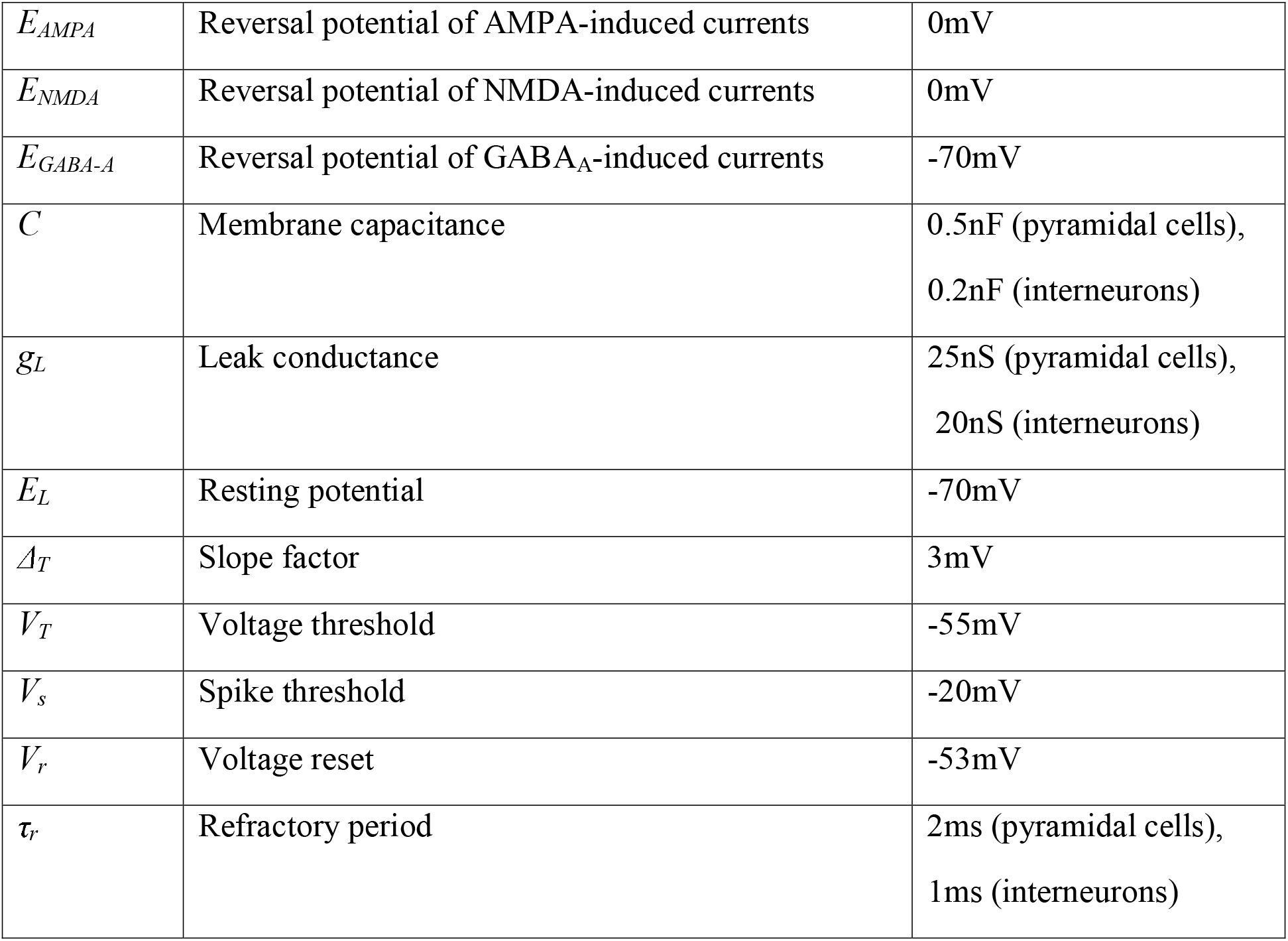
Parameter values for the neural model used in the simulations.

### Simulation of tDCS-Induced Currents

We simulated depolarizing tDCS by injecting a depolarizing transmembrane current into each pyramidal cell and hyperpolarizing current into each interneuron [15,36], and hyperpolarizing tDCS by adding hyperpolarizing current into pyramidal cells and depolarizing current into interneurons, a distinction which arises in cortex due to differences in orientation and cellular morphology. Note however that the results of our model are robust against these assumptions, and remain qualitatively similar when omitting current from interneurons (see below).

The simulated current was added to the input to each exponential LIF neuron:

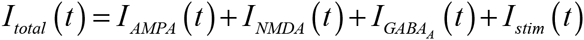

where *I_stim_(t)* is the tDCS current at time *t*.

The simulated tDCS current was applied for the duration of each stimulation trial, beginning at trial onset. Depolarizing tDCS was simulated by injecting 0.5pA into pyramidal cells and -0.25pA into interneurons, while during hyperpolarizing tDCS stimulation pyramidal cells were injected with -0.5pA and interneurons 0.25pA [15,36]. In additional simulations we applied stimulation only to the pyramidal populations; the results were qualitatively similar see Table S1-S6 and Discussion, [15]). The injected current simulating tDCS slightly changed the resting membrane potential of each neuron (±0.025mV with ±0.5pA injected current, ±0.0125mV with ±0.25pA), within the range found by *in vitro* tDCS studies [28,30,40].

### Simulation of the Perceptual Decision Making Task

One advantage of computational modeling is that the model can be run for many more trials than can feasibly be tested in human participants. However, this can lead to spuriously low-variance model predictions that cannot reliably be compared with human data. In order to fairly compare model and human behavioral performance, we generated 20 virtual subjects and assessed the effects of stimulation in each subject, under the same six coherence levels that human participants were tested with (0, 3.2, 6.4, 12.8, 25.6, and 51.2%), with 20 trials at each level. The sum of the two task-related inputs always equaled 80Hz (at coherence=0% both inputs were at 40Hz, and coherence=51.2% one input was at 60.48Hz and the other at 19.52Hz), so the total strength of the input received by the network remains equal across all conditions. Each trial lasted for 4s, with task-related input applied from 1-3s. This procedure was repeated for control (no stimulation), depolarizing, and hyperpolarizing stimulation conditions, respectively. In all stimulation conditions, stimulation was applied for the entire duration of the simulation: prior to and during the task-related inputs, thus matching the experimental procedure in humans where we applied tDCS during performance of the task.

### Human Participants

24 neurologically healthy volunteers participated in the experiment (7 male, aged 23.75±4.25). Participants gave their informed consent before participating and the local ethics committee approved the experiments (reference number 5833/001). Four participants were excluded from analysis due to poor performance on the task (accuracy threshold outliers as determined by the median deviation of the medians, [66]).

### Behavioral task

Participants completed a perceptual decision making task. Participants sat comfortably at a desk in front of a computer and responded to visual stimuli displayed on a screen by pressing keys on a keyboard. The screen had an update rate of 60Hz and was placed 76cm from the participants. On each trial, participants were required to fixate in the center of a screen. After 500ms a RDK was displayed and participants were required to press a key as soon as possible to indicate whether the direction of coherent motion was to the left or the right (Figure 4A). The RDK consisted of a 5° diameter circular aperture centered on the fixation point [47] with 0.1° diameter dots at a density of 16.7 dots/deg^2^/s [67], each moving at 5°/s [68]. The percentage of coherently moving dots was set randomly in each trial to 0, 3.2, 6.4, 12.8, 25.6, or 51.2%. Trials ended once a response had been made or after a maximum of 1s if no response was made. The inter-trial interval was 1-2s and varied depending on the response time of the previous trial to make all trials the same length. Participants were shown cumulative feedback at the end of each block displaying % correct, the mean response time in the most difficult trials, and # correct responses / minute. Before each session, participants completed a training block in which trial-by-trial feedback was given during the first ten trials. We used a within-subject design in which each participant completed three sessions (depolarizing stimulation; hyperpolarizing stimulation; no stimulation). The order of the stimulation conditions was balanced across participants.

### Transcranial Direct Current Stimulation

Transcranial Direct Current Stimulation (tDCS) was applied over the left dorsolateral prefrontal cortex (Figure 4B) using a battery-driven multi-channel direct current stimulator (NeuroConn, GmbH). The dlPFC was identified by MNI coordinates reported for activation within the superior frontal sulcus in posterior left dlPFC during perceptual decision making regardless of the response modality ([-23 29 37], [12]). We here optimized electrode positions for targeting of the left dlPFC using MRI-derived head models of electric field (EF) distributions to maximize current flow through this voxel (HD-Explore and HD-Targets software, v4.0, Soterix Medical, New York, NY, USA.). We determined that electrode positions 5cm medial and lateral to the nearest point to the dlPFC voxel on the scalp maximized current flow through the voxel (Figure 4B). Inward (relative to the cortical surface) currents should have an opposite effect on neural polarization to outward currents [28]. We therefore positioned the electrodes to generate depolarizing or hyperpolarizing currents in the left dlPFC [15,34]. Placing the reference electrode in the medial position maximized inward current and depolarization, while placing it in the lateral position maximized outward current and hyperpolarization [28,30,40,53,70].

Having identified the scalp-based markers for optimal stimulation of the left dlPFC, individualized electrode positions for each participant were derived using their structural MRI scan. Each participant’s was aligned to the MNI template and the dlPFC coordinate was localized in native space using the inverse co-registration transformation. The coordinate was then used in the neuronavigation software (Visor) to mark a target on the forehead corresponding to a location on the scalp radial from the coordinate. During depolarizing stimulation, the anode was positioned 5cm laterally from this target and the cathode was positioned 5cm medially. During hyperpolarizing stimulation the electrode polarities were reversed.

Participants completed a total of three sessions spaced approximately one week apart (Figure 4C). In the first session, participants completed three blocks of 200 trials each with three short breaks. Each block lasted 20min. During each session with stimulation (depolarizing or hyperpolarizing with the session order balanced across participants), participants completed two block of trials with sham stimulation, and one with depolarizing or hyperpolarizing stimulation. The first block was always sham and the order of the second and third blocks was balanced across participants. This controlled for possible learning effects by including a sham block directly preceding each stimulation block for comparison. During stimulation blocks, tDCS was applied for 20 min at 2 mA. During sham blocks the stimulation was ramped up to 2mA over 10s, stimulated for 30s, and then ramped down to 0 over 10s.

### Model and Task Implementation and Data Analysis

All of our model simulations were implemented in the Python programming language using the Brian simulator [71]. The human behavioral task was implemented in Python using PsychoPy [72]. Behavioral data of virtual and human participants and mean firing rates of the model were analyzed using SPSS (v21). We report two-way ANOVA tests for the main effects of stimulation condition (depolarizing, hyperpolarizing, and sham or no stimulation in the case of the model) and motion coherence as well as their interaction on each measured variable. The Greenhouse-Geisser correction of degrees of freedom was used when the sphericity assumption was violated (indicated by ε). Pairwise t-tests (Bonferroni corrected for multiple comparisons) were conducted between the stimulation conditions at each coherence level where a significant interaction effect was detected.

